# Protease Activity Profiling Via Programmable Phage Display

**DOI:** 10.1101/2020.05.11.089607

**Authors:** Gabriel D. Román-Meléndez, Thiagarajan Venkataraman, Daniel R. Monaco, H. Benjamin Larman

## Abstract

Endopeptidases catalyze the internal cleavage of proteins, playing pivotal roles in protein turnover, substrate maturation and the activation of signaling cascades. A broad range of biological functions in health and disease are controlled by proteases, yet assays to characterize their activities at proteomic scale do not yet exist. To address this unmet need, we have developed SEPARATE (<u>S</u>ensing <u>E</u>ndo<u>P</u>eptidase <u>A</u>ctivity via <u>R</u>elease and recapture using fl<u>A</u>nking <u>T</u>ag <u>E</u>pitopes), which uses monovalent phage display of the entire human proteome at 90-aa peptide resolution. We demonstrate that SEPARATE is compatible with several human proteases from distinct catalytic classes, including Caspase-1, ADAM17, and Thrombin. Both well-characterized and newly identified substrates of these enzymes were detected in the assay. SEPARATE was used to discover a non-canonical Caspase-1 substrate, the E3 ubiquitin ligase HUWE1, a key mediator of apoptotic cell death. SEPARATE is a novel methodology to enable efficient, unbiased assessment of endopeptidase activity using a phage-displayed proteome.

## Main

Proteases catalyze the irreversible hydrolysis of peptide bonds with consequences that include target destruction, protein maturation, and signal transduction. These enzymes participate in diverse biological functions, including tissue remodeling and morphogenesis, infection, blood coagulation, neoplasia, and cancer metastasis.^1–4^ Their enzymatic activities can therefore serve both as valuable diagnostic biomarkers and as therapeutic targets.^5–9^ There are 1,252 putative human proteases belonging to five families, accounting for ~3.5% of the human proteome.^10^ Given their importance and diversity, there is an unmet need for unbiased techniques to profile the activity of proteases, both in isolation and as components of complex biological mixtures. The physiological substrates of only a small fraction of proteases have been characterized in some detail; even for these enzymes, their full complement of substrates remains unknown.

Currently, unbiased protease profiling approaches are based on mass spectrometry^11,12^ and tend to be both cumbersome and expensive. Targeted activity-based profiling techniques^13,14^ can detect active proteases, but are typically limited by lower levels of assay multiplexing and are restricted to enzymes with well characterized substrates.^15,16^ Attempts to characterize cleavage motifs have also utilized the bacteriophage display of random peptide libraries^17–19^, but these types of analyses are typically difficult to interpret.

Here we present SEPARATE (<u>S</u>ensing <u>E</u>ndo<u>P</u>eptidase <u>A</u>ctivity via <u>R</u>elease and recapture using fl<u>A</u>nking <u>T</u>ag <u>E</u>pitopes), a highly multiplexed protease profiling platform that combines a complete human proteome library cloned into a novel T7 phage display vector and quantitative analysis via next generation DNA sequencing (NGS). SEPARATE enables unbiased, low-cost, and high sample throughput characterization of human protease activities, thereby overcoming key limitations of current approaches. ~250,000 oligonucleotides encoding 90 amino acid overlapping human peptide tiles, with 45 amino acid overlaps, covering the entire reference human proteome^20^, were synthesized and cloned as a pool into the T7-SEPARATE phage display vector (**Fig. 1A**). This vector displays a library peptide as a C-terminal fusion to the 10B T7 capsid protein, a distally flanking (C-terminal) biotinylated AviTag for library immobilization and a proximally flanking (N-terminal) 3x FLAG tag for recapture of released phage particles (**Fig. 1B**). The DNA sequences corresponding to the displayed peptides in the recaptured phage particles can then be amplified by PCR and quantified using NGS (**Fig. 1C**). SEPARATE detects proteolysis-dependent enrichments of each peptide in the recaptured library and is amenable to cost-reduction via sample multiplexing using barcodes introduced during PCR amplification.

**Figure 1.**
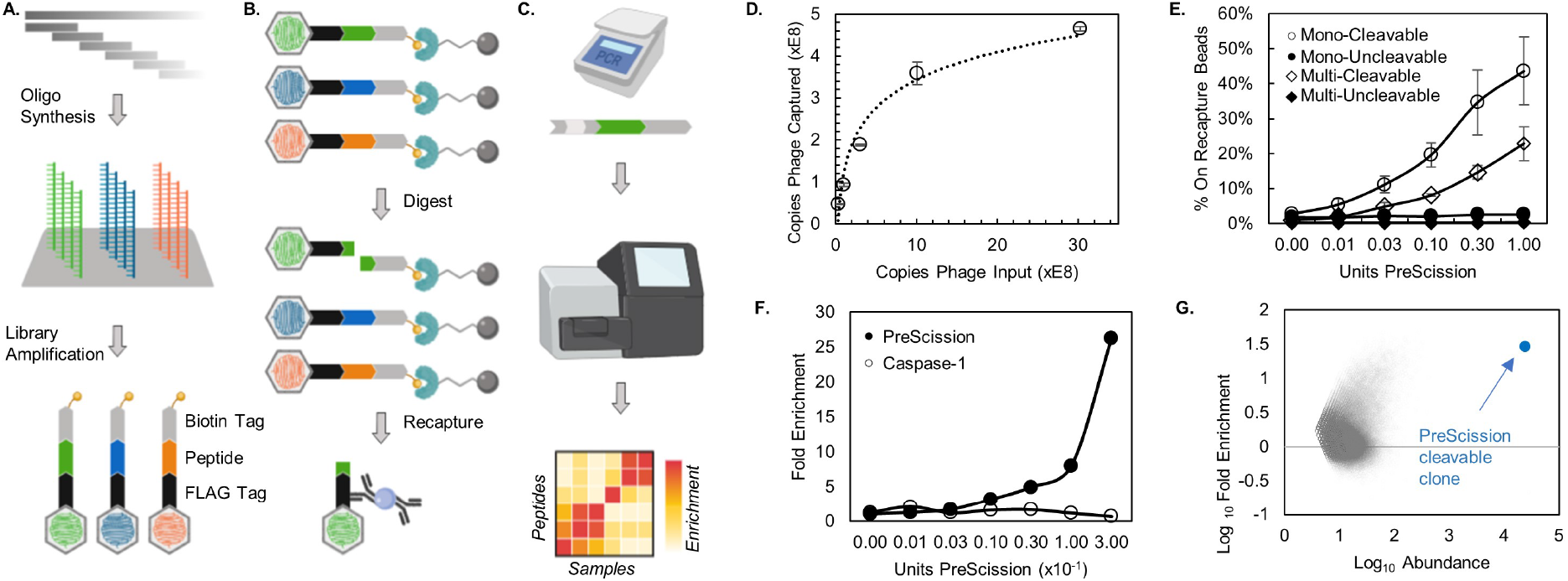
SEPARATE design and workflow. **(A)** An oligonucleotide library encoding >29,000 human protein isoforms as 45 amino acid overlapping 90-mer tiles is synthesized on a DNA microarray and cloned into the T7-SEPARTATE vector. This monovalent display vector flanks a library peptide by an N-terminal 3xFLAG tag and a C-terminal biotin labeling tag. **(B)** The biotin tagged library is immobilized on streptavidin-coupled magnetic beads, digested with a protease containing solution and digested phage clones are recaptured on M2 FLAG antibody coated protein G beads. **(C)** Recaptured phage clones are amplified by PCR and their clonal abundance quantified by deep sequencing to generate a fold-enrichment matrix. **(D)** T7-SEPARATE clones are biotin labeled in vivo and can be immobilized on streptavidin-coupled magnetic beads. **(E)** A T7-SEPARATE clone encoding the PreScission cleavage motif -LGVLPG/GP- was prepared using both the T7Select1-2b monovalent (○) and the T7Select1-3b multivalent (◇) T7 vector scaffolds. A negative control clone lacking the PreScission cleavage motif is not enriched on M2 FLAG antibody coated beads for both monovalent (●) and multivalent (◆) display scaffolds. **(F)** The PreScission digestible clone spiked into the human library demonstrates a dose-dependent enrichment (●); no enrichment is observed when digested with Caspase-1 (○). **(G)** MA plot analysis from a PreScission digest compares a peptide’s enrichment (Log[Protease/Buffer]) against its relative abundance in the human library (0.5*Log[Protease*Buffer]).

Irreversible library pre-immobilization has several advantages, including concentration of the library into small assay volumes, flexible buffer exchange for protease compatibility, and the removal of defective phage clones displaying peptides with frameshift or non-sense mutations. The C-terminal biotinylated AviTag provides near irreversible immobilization to streptavidin coated magnetic beads, which minimizes the amount of non-specific phage particle release and thus the background noise of the assay, even during lengthy digest reactions. Initial experiments revealed that the AviTag is sufficiently biotinylated during phage replication in *E. coli* cells to allow for robust binding to streptavidin beads. Immobilizing 10^8^ phage particles of the ~250,000-member human proteome library ensures that each peptide is represented greater than 100 times on average (**Fig. 1D**). Pre-blocking the streptavidin-coated beads with free biotin confirmed that ~99% of the bound phage are indeed immobilized via their C-terminal biotin tag (data not shown). Recapture of released phage particles via a proximal 3x FLAG tag reduces background noise from phage detached by physical dissociation of the displayed peptide, concentrates the recaptured phage into a small volume for PCR, while also removing potential PCR inhibitors present in the digest reaction.

We reasoned that monovalent peptide display would enhance detection sensitivity by requiring just a single protease cleavage event to release a target phage particle. To test this hypothesis, positive and negative controls were constructed, which could be monitored via quantitative real-time PCR. A cleavable 90-aa substrate for the commercially available PreScission enzyme (human rhinovirus 3C protease, GE Life Sciences, MA) served as a positive control, whereas a randomly selected 90-aa human peptide from the proteome library served as an uncleavable negative control. Both controls were also subcloned into a mid-copy (multivalent) version of the T7-SEPARATE vector, which displays 10B-fused peptides at a copy number between 5 and 15 per phage particle. At all concentrations of PreScission tested, the monovalent display format provided a substantial increase in the number of phage particles released and recaptured (up to 53% and 43%, respectively, of the total number of pre-immobilized phage particles), in comparison to the same conditions but using the multivalent display format (up to 24% and 22.5%, respectively; **Fig. 1E** and **Fig. S1**). When PreScission enzyme was omitted from the reaction (a ‘mock’ digest), only ~1% of the immobilized cleavable phage was detected on the recapture beads. In parallel, we measured the recapture of the uncleavable peptide, in which case ~1% of the immobilized phage particles were similarly recaptured, whether the PreScission enzyme was included in the digest reaction or not. Non-targeted peptides are therefore expected to contribute uniformly to the low background, both in the presence and absence of a test protease. Comparing the proportions of control peptide-displaying clones for the highest PreScission concentration versus the mock digest, the monovalent T7-SEPARATE format was found to perform at a signal-to-noise ratio of 23.4 versus 11.86 for the multivalent format.

To assess the performance of a proteomic-scale SEPARATE assay, the PreScission-cleavable clone was spiked into the complete human library at a ratio of 1 to 100. Recapture of the cleavable peptide demonstrated a protease concentration-dependent efficiency, with the highest concentration resulting in a 26-fold increase versus a mock digest or a non-PreScission protease (**Fig. 1F**). Furthermore, an MA-plot demonstrates that the PreScission-cleavable peptide displaying clone is indeed the most abundant clone on the anti-FLAG recapture beads and is also the most differentially enriched clone (**Fig. 1G**). Interestingly, a number of statistically significant candidate PreScission substrates were also identified in this experiment.

We next wondered whether SEPARATE could be used to identify novel and biologically relevant substrates, even for well-studied proteases. Caspase-1 plays a key effector function as part of the inflammasome complex by producing mature interleukin-1β (IL-1β)^21^ and activating pyroptosis via cleavage of gasdermin D.^22,23^ In our study, Caspase-1 was found to significantly cleave ~250 human peptide sequences corresponding to 230 unique genes. Peptides cleaved by Caspase-1 in the SEPARATE assay were analyzed using a motif detection algorithm, EpitopeFindr, which performs BLAST alignment of all peptides against each other to identify shared stretches of sequence homology.^24^ The results of this analysis were visualized as a network graph in which peptides were linked based on their alignments (**Fig. 2A**). Peptides within the largest cluster identified the multiple sequence alignment logo -DCXDXXDE- which strongly resembles “canonical” motifs reported for Caspase-3 and Caspase-8.^25^

**Figure 2.**
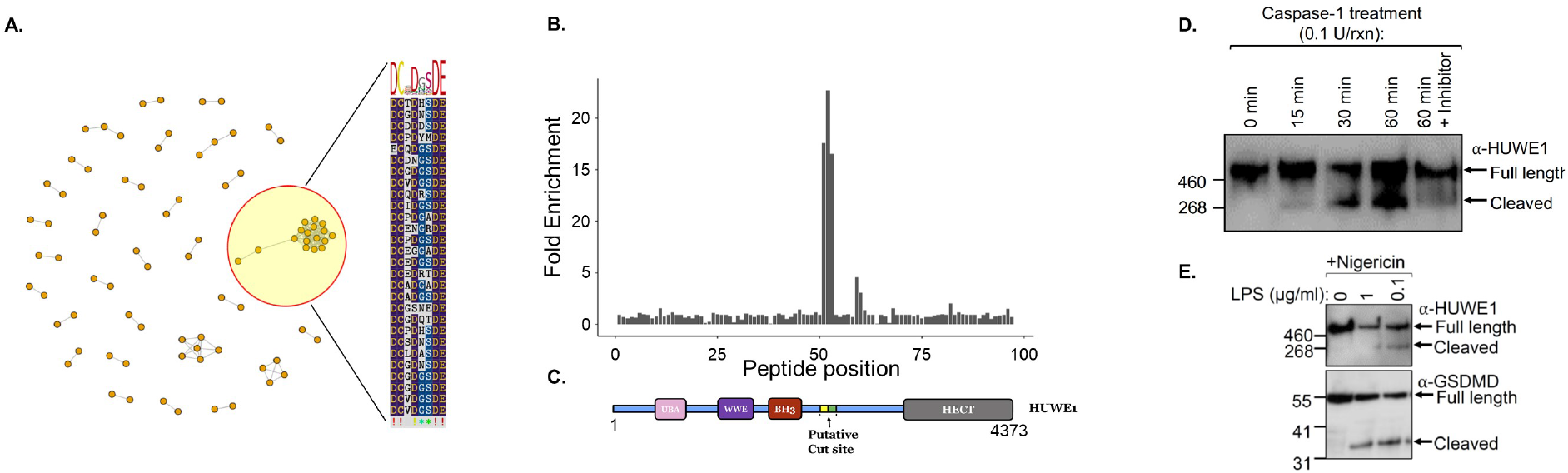
HUWE1 is a novel Caspase-1 target. **(A)** EpitopeFindr output as a network graph. Nodes represent Caspase-1 digested peptides and edges indicate regions of sequence homology. Multiple sequence alignment of the cluster containing the most peptides reveals a known Caspase cleavage motif. **(B)** HUWE1 peptide tiles 51, 52 and 53 are cleaved by Caspase-1. **(C)** Putative HUWE1 cleavage site identified by SEPARATE. **(D)** Western blot analysis of the time-dependent cleavage of HUWE1 in unstimulated THP-1 lysate after addition of recombinant Caspase-1. **(E)** HUWE1 (upper panel) and Gasdermin D (GSDMD, lower panel) are cleaved by endogenous Caspase-1 in THP-1 cells upon inflammasome activation with indicated amounts of LPS and 2.5mM Nigericin.

Interestingly, three peptides (two overlapping) derived from the protein HECT, UBA and WWE Domain Containing E3 Ubiquitin Protein Ligase 1 (HUWE1) were among the most significantly enriched, yet did not harbor the canonical cleavage motif (**Fig. 2B** and **Table S1**). HUWE1 (also known as LASU1, UREB1, MULE or ARF-BP1) is a well-known regulator of apoptosis, proliferation, DNA damage, and stress responses;^26–29^ Caspase-1 cleavage of HUWE1 has not been previously reported. The HUWE1 protein is 482 kDa; Caspase-1 cleavage is predicted to produce two fragments of ~250 and ~230 kDa (**Fig. 2C**). We sought to validate HUWE1 as a Caspase-1 substrate using THP-1 human monocytic cells, which are commonly used for studies of inflammasome activation. Addition of recombinant Caspase-1 to unstimulated THP-1 cell lysates resulted in robust cleavage of endogenous HUWE1 (**Fig. 2D**). Physiological inflammasome activation in THP-1 cells using LPS and Nigericin resulted in Caspase-1 activation and the cleavage of endogenous HUWE1 into fragments of the size predicted by SEPARATE (**Fig. 2E**). Considering the role of HUWE1 in promoting apoptotic cell death, we hypothesize that its destruction by Caspase-1 could tip the balance towards inflammation-associated pyroptosis. These results confirm that SEPARATE can be used to identify novel proteolytic substrates of putative biological consequence, even for well-characterized proteases.

For SEPARATE to be useful in the analysis of complex cell or tissue lysates, it must have sufficient sensitivity at physiologically relevant enzyme concentrations. To this end, we performed SEPARATE on the human library and specifically focused on previously known substrates of well-characterized recombinant proteases from diverse catalytic classes. Testing serial dilutions of Caspase-1 (a cysteine protease), Thrombin (a serine protease), and Adam17 (a zinc-dependent metalloprotease), concentration-dependent enrichments of known substrates were indeed detected for each enzyme at or below their reported physiological concentrations (**Fig. 3A**); these substrates ranked among the most significant enrichments for each of the corresponding enzymes (**Fig. 3B-D**). Additionally, to assess whether SEPARATE could be performed on a complex biological mixture, we used unstimulated THP-1 cell lysate supplemented with 0.1 units of recombinant Caspase-1. In this setting, HUWE1 peptides were again among the most significantly enriched targets (**Fig. 3D**).

**Figure 3.**
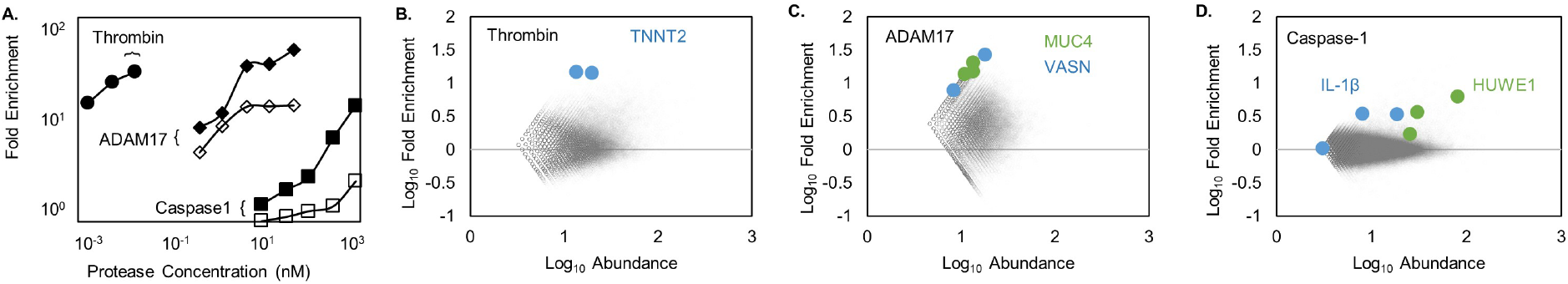
SEPARATE detects known protease targets at physiological concentrations. **(A)** Aggregated data from series of digests using Thrombin, ADAM17 and Caspase-1. Physiological concentrations for Thrombin and ADAM17 are 1-1,000 nM and 1-10 nM; A range for Capsase-1 is not reported. Cardiac troponin T (TNNT2; ●) is a known Thrombin substrate; Mucin 4 (MUC4; ◆) and Vasorin (VASN; ◇) are known substrates of ADAM17; IL-1β (□) and HUWE1 (■) known substrates of Caspase-1. **(B-C)** MA plot analysis of Thrombin (2 pM), ADAM17 (0.4 nM) and **(D)** Caspase-1 in THP1 lysate (0.3 μM).

The performance of the SEPARATE assay is expected to depend on protease-substrate requirements as well as biophysical features of the peptide library. We have verified by PCR that 90% of the immobilized human library lacks in-frame mutations. However, the well characterized Caspase-1 substrate Gasdermin-D did not score in the assay, and this was true for several other canonical enzyme-substrate pairs. Using a 56-amino acid mouse peptidome library in which the Gasdermin-D cleavage peptide is better represented, robust Caspase-1 dependent cleavage was indeed observed (**Table S2**). In general, poor library representation of some peptides, particular enzymes’ requirements for substrate conformational structure, cofactors, and/or post-translational modifications, are all expected to cause false negative SEPARATE assay results.

Currently, there are large gaps in our knowledge of protease substrates, even for well-studied enzymes like Caspase-1. We have therefore developed the SEPARATE system for unbiased, high throughput, inexpensive, and automatable protease activity profiling using peptidome libraries monovalently displayed on phage. While not always able to localize precise cleavage sites, SEPARATE can identify novel substrates and narrows down the recognition motif to 45-90 amino acids. In a proof-of-concept study, we detected peptides that together recapitulated a canonical Caspase motif and identified a novel, non-canonical, physiological substrate, HUWE1, which was validated using standard approaches. The SEPARATE methodology is broadly applicable as it can be readily adapted to any soluble endopeptidase of interest, including endogenous and pathogen-associated proteases present within complex biological samples. We therefore expect that unbiased protease profiling via approaches like SEPARATE will identify valuable new disease biomarkers and therapeutic targets.

## Materials and Methods

### Materials

All materials not described below were purchased from commercial suppliers and were of the highest grade available. Caspase-1, ADAM17, Thrombin, and PreScission proteases were purchased from BioVision, RnD Systems, SignalChem, and GE Life Sciences respectively.

### Construction of T7 SEPARATE Phage Vector

The SEPARATE vector (T7-SEPARATE) was constructed by cloning a custom-designed gBlock gene fragment (IDT) into the BamHI/SalI site of the low-copy T7Select1-2b phage vector (Millipore). The gBlock fragment is based on a 90-amino acid β-lactoglobulin peptide sequence containing two PreScission protease cleavage sites (LEVLFQGP).^30^ C-terminal to the peptide is a V5 tag, followed by a TEV cleavage sequence, and then an AviTag^31^ biotinylation sequence (GLNDIFEAQKIEWHE), which is enzymatically conjugated with a single biotin moiety *in vivo* during phage replication in *E. coli*. N-terminal to the displayed peptide is a 3x-FLAG tag followed by an enterokinase cleavage site. Restriction sites EcoRI/XhoI were placed upstream and downstream of the engineered β-lactoglobulin peptide to allow for library subcloning into the T7-SEPARATE vector.

### Subcloning the Human Peptidome Library into the T7-SEPARATE Vector

T7-Pep2,^32^ a complete human proteome library was restriction cloned into the T7-SEPARATE vector using the EcoRI/XhoI sites.^32,33^ The library was packaged *in vitro* using the T7Select Packaging Kit (Millipore Sigma) and expanded by plate lysate amplification to obtain an average clonal representation of ~100 plaques per peptide. To select for AviTag-displaying peptides, the initial expansion of this library was immobilized on streptavidin beads, washed to remove unbound phage particles (to remove prematurely truncated or out-of-frame peptides which will lack the downstream AviTag), digested with enterokinase, and re-expanded by plate lysate amplification at >100-times representation per peptide. Bacterial debris and large particulates were removed by centrifugation, followed by filtration through a 0.22 μm PES membrane, and the clarified library pool was stored at −80°C in 10% DMSO. The quality of the library peptides was assessed by Sanger sequencing of individual plaques from the packaging expansion and the clonal distribution assessed by Illumina sequencing.

### SEPARATE Assay: Immobilization of the T7-SEPARATE Library

Streptavidin-coupled magnetic beads (Dynabeads M-280 Streptavidin, ThermoFisher Scientific) were washed (TBS, pH 7.4, 0.01% NP40) and resuspended in binding buffer (TBS, pH 7.4, 0.001% NP40) containing 2E9 plaque forming units per 10 μl of bead slurry. End-over-end mixing at room temperature was performed for one hour. Phage-coated beads were washed three times with wash buffer and transferred into two protein LoBind Eppendorf tubes blocked with TBS, pH 7.4 containing 1% BSA.

### SEPARATE Assay: Protease Digestion

Protease digestions took place in 50 μl of appropriate buffer alone or containing the protease. Digestion occurred overnight at room temperature with end-over-end rotation. Digest reactions were quenched by addition of 100 μl TBS, pH 7.4, 0.01% NP40 containing AEBSF (2 mM), Aprotinin (0.3 μM), Bestatin (116 μM), E-64 (14 μM), Leupeptin (1 μM), and EDTA (1 mM) for thirty minutes at room temperature. To remove any residual biotinylated phage that may have been nonspecifically released, the 150 μl supernatant of the quenched digest reaction is then incubated with a fresh 10 μl of slurry volume M280 streptavidin beads (pre-washed three times with wash buffer) in a LoBind Eppendorf tube blocked with TBS, pH 7.4 containing 1% BSA at room temperature for one hour with end-over-end rotation.

### SEPARATE Assay: Recapture of Cleaved Phage Clones

10 μl slurry volume of protein G magnetic dynabeads were washed in buffer (TBS, pH 7.4, 0.01% NP40) and then resuspended in 50 μl TBS, pH 7.4, 0.01% NP40 containing 4 μg of M2 FLAG antibody for thirty minutes at room temperature with end-over-end rotation. The 150 μl quenched digest reaction is added to the 10 μl FLAG-coated Protein G beads and incubated for 60 minutes at room temperature with end-over-end rotation to capture phage particles released at the protease digest step. Beads are rinsed once with 100 μl wash buffer and stored at −80°C until PCR amplification.

### SEPARATE Assay: Amplification and Sequencing of Recaptured Phage Clones

Library PCR preparation, high-throughput DNA sequencing, and peptide read count data generation was performed as described previously.^32^ Briefly, library peptide inserts are amplified by resuspending 10 μl of FLAG-coated Protein G beads in PCR1 master mix containing the T7-Pep2_PCR1_F forward primer (ATAAAGGTGAGGGTAATGTC) and a T7-SEPARATE vector specific reverse primer (CTGGAGTTCAGACGTGTGCTCTTCCGATCAACCCCTCAAGACCCGTTTA), which includes an adapter sequence for sample-specific barcoding and Illumina P7 adapter incorporation during a subsequent PCR2 reaction. The PCR2 amplicons are pooled and sequenced using an Illumina NextSeq instrument in standard output mode to obtain single-end 50 nucleotide reads. Dual indexed sample demultiplexing and clonal read count determination were performed using exact sequence matching. Read counts were normalized using a ‘random peptide normalization’ method, which attempts to make data comparable between samples by calculating a normalization factor based on ‘background’ recaptured phage clones. To calculate the normalization factor, 100 peptides were randomly selected from the mock digest conditions with a read count ranging between 10 and 40. The median read count value for these 100 peptides is calculated for each sample. The random peptide selection and median calculation is performed 20 times and the average of the 20 median values is calculated for each sample. Finally, the normalization factor is calculated by dividing the average median value for each sample by that for one of the mock digest conditions. Read counts are then converted to normalized read counts by dividing each sample’s read count values by the normalization factor. Fold enrichments are calculated for each peptide by dividing their normalized read counts in the digest condition by the normalized read counts in the mock digest condition. The fold enrichments can be visualized in an MA plot by transforming the normalized read count data into a log ratio (M, on the y-axis) and a mean average (A, on the x-axis) between the digest and mock digest conditions.

### THP1 cell culture

THP-1 cells were cultured in RPMI-1640 media with Glutamax (Gibco, ThermoFisher, Cat #61870127), supplemented with 10% FBS (Hyclone SH30071, GE Life Sciences), and 1X Antibiotic-Antimycotic (Gibco, ThermoFisher, Cat #15240062), referred to as complete RPMI. To differentiate THP-1 cells into macrophages, 1×10^6^ THP-1 cells were added per well of a 6-well plate in complete RPMI medium containing 50 ng/ml PMA (diluted in RPMI medium from a 1 mg/ml PMA stock in DMSO). After 48 hours when the cells become adherent, the medium with PMA was removed, cells washed in 1X PBS once, and fresh complete RPMI was added. Cells were allowed to rest for an additional 24 hours before inflammasome activation with LPS and Nigericin

## Acknowledgements

This work was made possible by generous support from The Sol Goldman Pancreatic Cancer Research Center and the Emerson Collective Cancer Research Fund. We are grateful to Stephen Elledge (Harvard Medical School) for generously providing the T7-Pep2 90-mer human peptidome library. Uri Laserson (Mount Sinai Icahn School of Medicine) designed the 56-mer mouse peptidome library. Thanks to Dennis Grab for helpful discussions.

## Conflicts of Interest

G.R-M., T.V. and H.B.L. are listed as inventors on a patent application describing the SEPARATE assay.

## Supplementary Materials

**Supplemental Figure 1.**
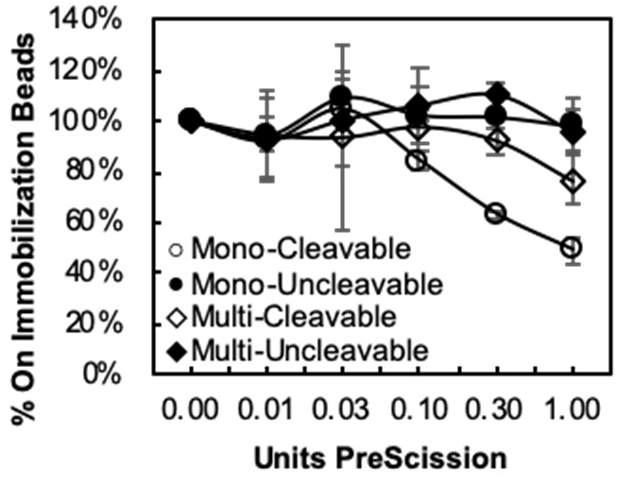
A T7-SEPARATE clone encoding the PreScission cleavage motif -LGVLPG/GP- was prepared using both the T7Select1-2b monovalent (○) and the T7Select10-3b multivalent (◇) T7 display vectors. The monovalent display results in a higher depletion of phage clones from the immobilization beads indicating a higher fraction is digested at a given concentration of PreScission protease. A negative control clone lacking the PreScission cleavage motif is not digested off of streptavidin-coated beads for both monovalent (●) and multivalent (◆) display vectors.

**Supplemental Table 1.** Top 20 enriched peptides from human library digestion with Caspase-1.

| Gene Name | Amino acid position | Caspase1 p_val | Caspase1 fold change |
| --- | --- | --- | --- |
| HUWE1 | 2296-2385 | 2.43E-04 | 22.71 |
| Efs | 181-270 | 1.34E-03 | 21.06 |
| LYST | 1171-1260 | 1.06E-03 | 21.04 |
| ERO1L | 91-180 | 6.53E-04 | 19.8 |
| Hyls1 | 1-90 | 9.59E-04 | 19.74 |
| znf462 | 1-90 | 3.89E-04 | 19.1 |
| BSN | 631-720 | 9.57E-04 | 18.78 |
| unmapped_NM | 91-180 | 7.81E-04 | 18.72 |
| TNS3 | 451-540 | 4.20E-04 | 18.67 |
| C14orf93 | 91-180 | 6.86E-04 | 18.22 |
| HUWE1 | 2251-2340 | 1.51E-03 | 17.58 |
| hoxa2 | 271-360 | 5.52E-04 | 17.06 |
| ZNF469 | 811-900 | 9.14E-04 | 16.76 |
| HUWE1 | 2341-2430 | 1.31E-03 | 16.55 |
| MAP3K14 | 496-585 | 1.42E-03 | 16.5 |
| HMGXB4 | 226-315 | 6.29E-04 | 16.49 |
| LOC100287737 | 496-585 | 1.70E-03 | 16.43 |
| UHRF1BP1L | 991-1080 | 3.04E-03 | 16.39 |
| ANAPC1 | 1-90 | 7.69E-04 | 16.35 |
| Hmcn2 | 1576-1665 | 6.90E-04 | 16.27 |

**Supplemental Table 2.**
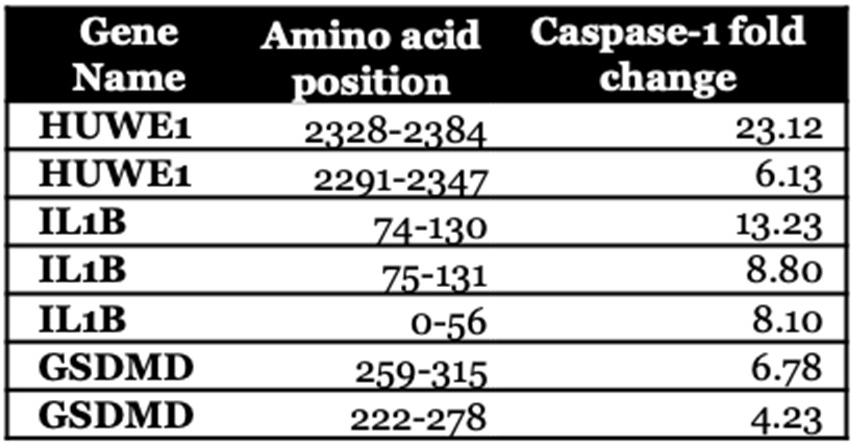
Selected peptides from mouse library digested with Caspase-1.

| Gene Name | Amino acid position | Caspase-1 fold change |
| --- | --- | --- |
| HUWE1 | 2328-2384 | 23.12 |
| HUWE1 | 2291-2347 | 6.13 |
| IL1B | 74-130 | 13.23 |
| IL1B | 75-131 | 8.80 |
| IL1B | 0-56 | 8.10 |
| GSDMD | 259-315 | 6.78 |
| GSDMD | 222-278 | 4.23 |

